# Heme Detoxification in the Malaria Parasite *Plasmodium falciparum*: A Time-Dependent Basal-Level Analysis

**DOI:** 10.1101/2025.03.06.641703

**Authors:** Larnelle F. Garnie, Timothy J. Egan, Kathryn J. Wicht

## Abstract

Malaria is a deadly disease for which therapeutic options are threatened by the rise of antimalarial resistance. Inhibiting the formation of hemozoin (the product of heme detoxification) in the digestive vacuole (DV) is the mechanism of action of numerous antimalarial drugs, including those in development as new therapies. This drug target remains attractive as hemozoin is an abiotic and non-mutable molecule, unique to the parasite. The underlying parasite biology of the heme detoxification pathway is complex and requires a deeper understanding. This study focuses on the DV of *Plasmodium falciparum*, utilizing confocal microscopy, immunoblotting and cellular fractionation techniques to study its native state over time. Using parameters such as the uptake into and growth of the DV, relative abundance of plasmepsins (PMs) I and IV and basal levels of hemoglobin, heme and hemozoin, it was found that DV physiology in chloroquine (CQ)-sensitive NF54 parasites follows three distinct developmental phases: the lag-type growth (20 to 28 h), rapid growth phase (28 to 40 h) and the plateau (40 to 48 h). These phases hold specific characteristics with respect to the investigated parameters. In addition, key differences between CQ-sensitive NF54 and CQ-resistant Dd2 parasites were observed.

## Introduction

*Plasmodium falciparum* parasites cause the most virulent form of malaria, a disease that infected 263 million people in 2023 and resulted in 597 000 fatalities. Strikingly, 94% of these cases and fatalities occurred in the World Health Organization (WHO) African Region.^1^ Currently, the WHO-recommended treatment for uncomplicated blood-stage malaria is the use of artemisinin-based combination therapies (ACTs), in which a fast-acting artemisinin derivative is administered in combination with a slow acting partner compound.^1–3^ However, concerns are rising over the continued efficacy of ACTs, owing to reports of partial artemisinin resistance in East Africa.^4^

Some of the widely used ACT combinations such as artesunate-amodiaquine, artesunate-mefloquine, artesunate-pyronaridine, artemether-lumefantrine, and dihydroartemisinin-piperaquine contain partner drugs that target essential processes within the heme detoxification pathway of the parasite.^2, 3^ Furthermore, many recent studies have identified new classes of antiplasmodium compounds that therapeutically act on this pathway.^5–10^

During the asexual blood stage (ABS), the symptomatic stage of a *P. falciparum* infection, parasites ingest and degrade hemoglobin resulting in the release of cytotoxic heme, which is subsequently detoxified through incorporation into a crystal called hemozoin. Inhibiting hemozoin formation remains an attractive approach, in spite of wide-spread drug resistance to the classical hemozoin formation inhibitor, chloroquine (CQ), given that the resistance mechanism of CQ involves drug-specific efflux from the DV by the chloroquine resistance transporter, *Pf*CRT. Notably, no changes in the process of hemozoin formation itself have been observed in strains resistant to hemozoin formation inhibitors. Indeed, many hemozoin formation inhibitors are active against the CQ-resistant (CQR) *Pf* strain, Dd2, which harbours eight mutations in *Pf*CRT compared with the wild-type CQ-sensitive (CQS) NF54 strain.^11–13^

Previous studies have shown that hemoglobin uptake commences during the ring stage through the formation of cytostomal invaginations containing hemoglobin.^14–18^ It has been suggested that these hemoglobin-filled vesicles then coalesce to form the central DV compartment.^16^ Although this is the most widely accepted uptake route, other uptake mechanisms such as the “Big Gulp” have also been suggested.^18^ Hemoglobin is then digested by a cohort of proteases localized to the DV.^19–22^ The aspartic proteases plasmepsin (PM) I, II, III (also known as histoaspartic protease) and IV, as well as cysteine proteases falcipain 2 and 3, have been shown to be involved in the initial catabolism of hemoglobin^19, 20, 23–27^ to produce oligopeptides resulting in the release of the heme moiety.^28^ The oligopeptides are further degraded by action of falcilysin, a zinc metalloprotein,^29^ dipeptidyl aminopeptidase 1 (DPAP 1),^30, 31^ and various aminopeptidases.^32–34^ The cytotoxic heme released through this catabolic process is then oxidized and incorporated into insoluble hemozoin crystals. This process is proposed to be lipid-mediated, although there may also be proteins, such as heme detoxification protein and lipocalin, involved.^35^ Interestingly, only 16% of amino acids produced from hemoglobin catabolism are used by the parasite.^36^ Instead, it has been postulated that hemoglobin is catabolized to create space for the parasite to grow and to maintain osmotic pressure within the RBC to prevent premature lysis.^37, 38^ Antimalarials including CQ, amodiaquine, mefloquine, pyronaridine, piperaquine and lumefantrine have been shown to inhibit β-hematin (synthetic hemozoin) formation in extracellular assays, whereas only a subset of these have demonstrated inhibition of hemozoin formation in cell-based assays. For example, true inhibitors of hemozoin formation cause a dose-dependent increase in free heme in treated parasites with a corresponding decrease in hemozoin in the cell fractionation assay.^11, 39, 40^ On the other hand, some compounds such as mefloquine, only show a decrease in hemozoin without an increase in heme. This effect may result from the inhibition of hemoglobin endocytosis, leading to a reduction in the total amount of released heme and therefore, hemozoin.^41^

These observations highlight the importance of whole-cell studies to inform the mode of action for compounds exerting their therapeutic effect within the DV. Furthermore, although specific steps within the heme detoxification pathway have been studied in the context of specific enzymes or drugs, a holistic view of the basal parasite biology underlying this pathway is required to better describe DV processes. For example, changes in the DV lumen volume, PM protein concentrations, and the basal-levels of heme-containing species during ABS parasite growth have not been extensively probed.

Thus, here we show changes in the DV lumen growth, excluding the hemozoin crystals, over time in CQ-sensitive NF54 and CQ-resistant Dd2 strains using confocal microscopy, and a macromolecular probe called pHrodo™ red dextran. This probe fluoresces only at low pH and was used to selectively stain the acidic DV.^42–44^ This approach allowed for live-cell imaging, which requires minimal pre-processing of the samples and largely maintains cellular integrity. To our knowledge this is the first use of pHrodo™ in this context. Three parameters: DV lumen volume, uptake into the DV and concentration of pHrodo™ within the DV were measured. In addition, we examined the wild-type PM I and PM IV protein levels over the trophozoite to schizont life stage using immunoblotting and densitometry techniques. Finally, we show differences in the basal levels of hemoglobin, heme and hemozoin between the NF54 and Dd2 strains using a heme fractionation assay.^39^ A culmination of these parameters allows for a more holistic overview of the DV and heme detoxification pathway.

## Results

### Digestive vacuole studies

#### Volume

Culture clock standards for NF54 and Dd2 were constructed over a consecutive 24-hour period. The cell count-normalized fluorescence channel 1 area (FL1-A), representative of SYBR™ Green, correlates with the DNA content which increases as the parasite ages. For all time-sensitive assays, the FL1-A standard histograms were overlaid with the experiment histograms, and the Giemsa-stain thin films compared to confirm parasite age (Figure **1**, Extended Data Figures 1 and 2).

**Figure 1.**
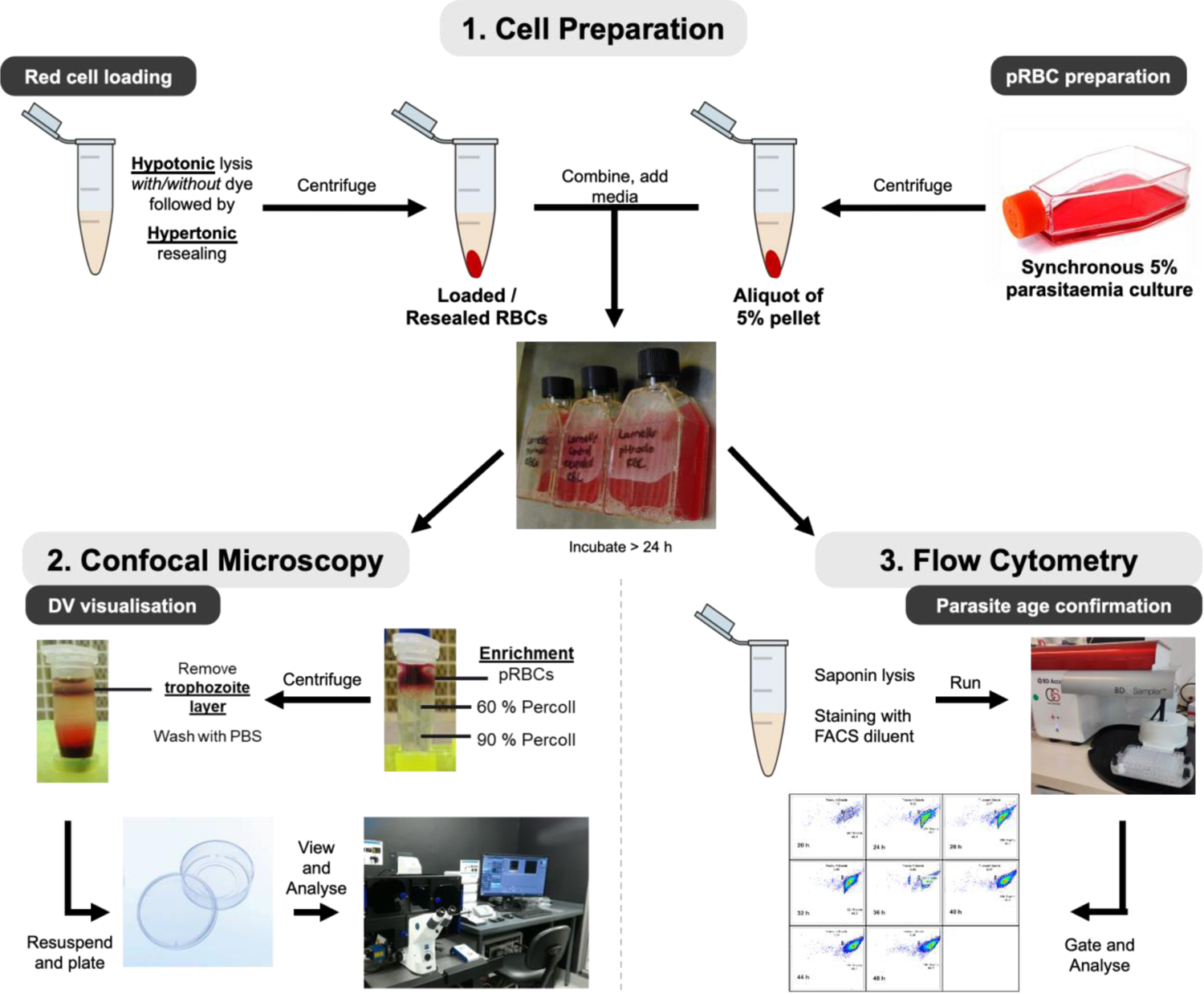
Method summary of confocal imaging and flow cytometry for NF54 parasites used to determine the volume of and uptake into the DV lumen. **1.** Cell preparation involved hypotonic lysis and hypertonic resealing of RBCs to produce resealed RBCs (with and without pHrodo). In addition, an aliquot of a 5% parasitemia culture was added to the resealed RBCs and media, followed by incubation at 37 °C for more than 24 hours. **2.** Purified trophozoite-containing pRBCs were obtained through enrichment, resuspended, and plated in imaging dishes. Images were then acquired using confocal microscopy. **3.** An aliquot of the pRBCs was saponin lysed, fixed, and stained in preparation for flow cytometry. The samples were run on a flow cytometer and the resultant populations gated and analyzed to confirm parasite age.

Red blood cell (RBC) ghosts were pre-loaded with pHrodo™ using hypotonic lysis and hypertonic resealing (Figure **1**). Parasites were then allowed to reinvade the loaded RBCs. Following the incubation period, Z-stacks of individual parasites were acquired on a confocal microscope (Figure **1**, Figure **2A** and Extended Data Figure 3) and these were analyzed in ImageJ to calculate the DV lumen volume (Figure **2B**).^45, 46^ Time 0/48 h and 0/46 h for NF54 and Dd2, respectively, were treated as the point at which the schizonts burst to release merozoites. At these time points, the DV lumen no longer exists as shown in the schizont inset of Figure **2B**, where the hemozoin is tightly wrapped in the DV membrane. Therefore, on this basis, the curve was extrapolated to the origin for both strains and to (48,0) and (46,0) for NF54 and Dd2, respectively.

**Figure 2.**
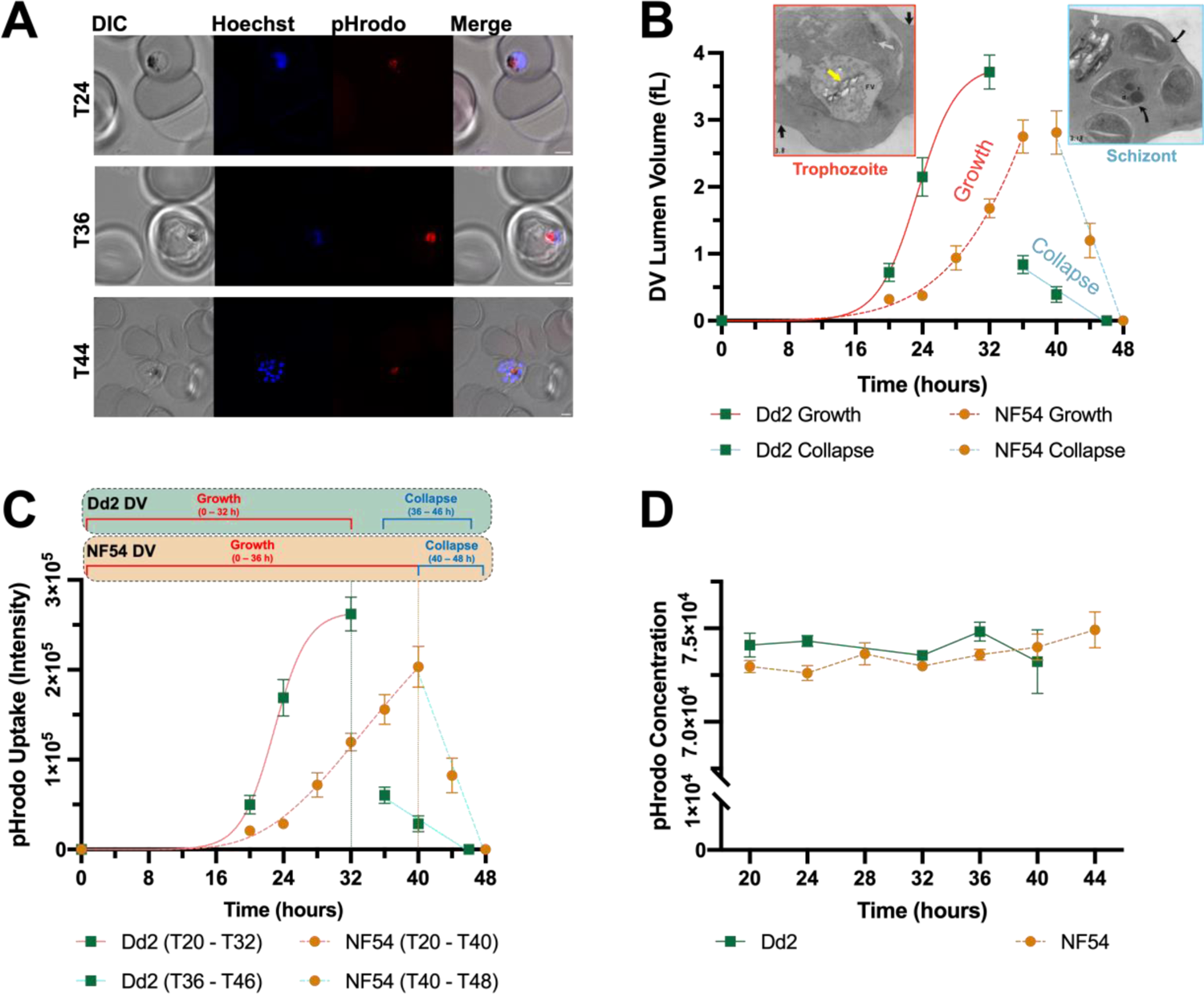
Digestive vacuole lumen volumetric and uptake analysis in *Pf* parasites (Dd2 and NF54) using confocal microscopy. Parasites were allowed to re-invade RBCs pre-loaded with pHrodo™-dextran red and imaged using the Z-stack functionality of an LSM 880 Airyscan confocal microscope (Zeiss). **A.** A single Z-stack slice of representative individual RBCs infected with NF54 parasites at various timepoints (stipulated on image). Hoechst (blue) stains the DNA while pHrodo™ (red) selectively stains the DV. All scale bars represent 2 µm. **B.** Growth of the DV lumen in *Pf* parasites as determined using the 3D Objects Counter (3D OC) calculation in ImageJ.^45, 46, 49^ Growth and collapse curves obtained for both strains were plotted on the same axis. Gompertz growth curves were fitted to the growth of the lumen and exhibited an r^2^ > 0.99 for both strains, while linear regression analysis of the collapse exhibited an r^2^ > 0.98 for both strains. Insets. TEM image of a trophozoite, magnification ×32 500, DV is labelled as FV (food vacuole). Yellow arrow = hemozoin crystals, black arrow = knobs on RBC membrane, white arrow = dense granules. TEM image of a late schizont stage, magnification ×32 500. Black arrows = separated merozoites, white arrow = redundant DV in which the hemozoin is “shrink-wrapped” in the DV membrane and devoid of any lumen space, r = rhoptries, d = dense granules. Images produced from unpublished data taken from Honors dissertation of Dr Joanne Egan with permission. **C.** The uptake of pHrodo™ into the DV lumen of *Pf* parasites measured by absolute fluorescence intensity using the “Analyze Particles” (AP) function in ImageJ. The phases of DV lumen growth and collapse are shown on the graphs for reference and the endpoint of the DV lumen growth is shown as a dotted line. NF54 uptake data was fitted to a Gompertz growth curve with r^2^ > 0.99 for 0 ≤ x ≤ 40 and a linear regression with r^2^ > 0.98 for 40 ≤ x ≤ 48. Dd2 uptake data was fitted to a sigmoidal growth curve (variable slope) with r^2^ > 0.99 for 0 ≤ x ≤ 32 and a linear regression with r^2^ > 0.97 for 36 ≤ x ≤ 46. **D.** The concentration of pHrodo™ within the DV in arbitrary units, calculated as intensity divided by DV lumen volume. For each data point in graphs n ≥ 15 (error bars indicate SEM).

The growth of the DV lumen between 0 and 36 h in NF54 most accurately fit a Gompertz growth curve (Figure 2B).^47, 48^ The plateau of the curve within the NF54 dataset was not reached before the DV lumen began to collapse at 40 h. Between 40 and 48 h, consistent with the appearance of schizonts, the DV lumen collapsed following a linear regression. In Dd2, the life cycle more accurately followed a 46-hour cycle, and the DV lumen expanded between 0 and 32 h, also according to the Gompertz growth curve. In this case, contrary to what was observed for NF54, the plateau was seen to begin near 32 h before the Dd2 DV lumen collapsed linearly between 36 and 46 h.

#### Uptake and concentration

In addition to studying the DV lumen volume, pHrodo™ was used to approximate the uptake into the DV via endocytosis. pHrodo™ was used as a proxy for hemoglobin on the basis that during the hypotonic lysis and hypertonic resealing of the RBCs, pHrodo™ was loaded into the RBC cytoplasm and was endocytosed as the parasite ingested the RBC cytosol. Using the “Analyze Particles” function in ImageJ,^45, 46^ the background-corrected, integrated density over each Z-stack was calculated and averaged over all the samples for each timepoint to determine the fluorescence intensity (Figure 2C) within the DV.

It was found that the data trends for uptake into the lumen closely resemble those of DV lumen growth. Between 0 and 40 h in NF54, the uptake into the DV was shown to also follow Gompertz growth, corresponding with DV lumen growth (Figure 2C). Furthermore, the collapse of the DV lumen between 40 and 48 h corresponds to a linear decrease in the amount of RBC cytosol taken up into the lumen. In contrast to the Gompertz growth curve fit to NF54 uptake, for Dd2 (Figure 2C), a more flexible sigmoidal curve (variable slope) was fitted to the DV uptake between 0 and 32 hours. Between 36 and 46 hours there was a linear decrease in the uptake into the lumen. The pHrodo™ concentration, calculated as the uptake averages divided by lumen volume, was found to be statistically constant over all timepoints in both NF54 and Dd2 (Figure 2D).

### PM I and IV protein analysis

#### Protein levels over time

Given the importance of the PMs in hemoglobin catabolism, PM I and PM IV were chosen as model enzymes for this study. It has been shown that these increase in abundance in aging parasites,^50^ therefore the time-dependent levels in NF54 parasites, as a representative model strain, were examined in this study. As an internal control, the constitutively expressed endoplasmic reticulum chaperone binding immunoglobulin protein (BiP), was used as a proxy for changes in the total protein over the trophozoite to schizont stage.^51, 52^ Immunoblots of parasite cell lysates were separated on 12% SDS-page gels, transferred to PVDF membranes and probed with anti-PM I, anti-PM IV and anti-BiP antibodies (Figure 3 and Extended Data Figures 4A-B). Dd2 PM I and PM IV knockout (KO) lines were used as controls to confirm specificity of the antibodies.^53^ Densitometric analysis of the protein levels per 50 000 parasites provided relative quantification of the three proteins and enabled the calculation of the relative amount of PM I and PM IV from 20 to 44 h, in NF54 parasites (Figures 3A-B). For both PM I and PM IV the calculated protein levels between 20 and 28 h remained at a similar level, after which there was a statistically significant increase from 28 to 32 h for both proteins. Between 32 and 44 h there was a gradual increase between consecutive timepoints with the levels reaching a plateau between 40 and 44 h.

**Figure 3.**
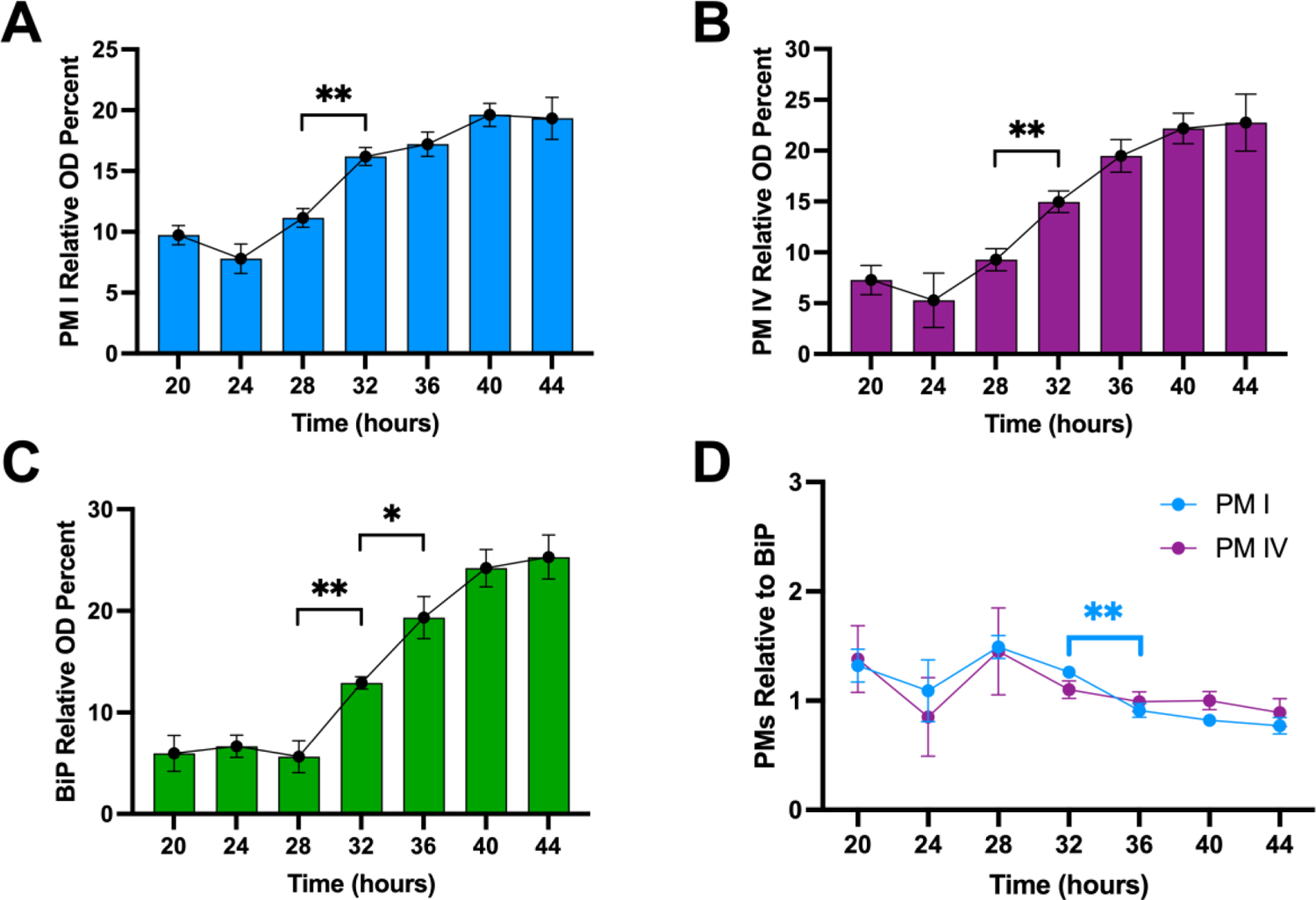
Cell lysates obtained from ∼50 000 parasites (as determined by flow cytometry) over the trophozoite to schizont phase were separated on a 12% SDS page and then transferred to a PVDF membrane. The membranes were probed with PM I (anti-rabbit, 1:10 000), PM IV (anti-mouse, 1:5 000) and BiP (anti-rabbit, 1:10 000) antibodies. Following chemiluminescent detection of the immunoblot signals, densitometric analysis of the blots were carried out using ImageJ as previously described.^54^ **A**., **B**. and **C**. show the relative amounts of PM I, PM IV and BiP, respectively. The y-axes are represented as relative OD percentages in arbitrary units. **D**. Relative amounts of PM I and PM IV normalized to BiP. Significance levels were calculated using two-tailed, unpaired student’s t-tests where N = 4 for each timepoint (* p < 0.05; ** p < 0.01, error bars indicate SEM).

The levels of BiP showed an overall increase from 20 to 44 h (Figure 3C) with the levels between 20 and 28 h remaining steady. The increase from 28 to 44 h, however, followed a steeper trend than PM I and PM IV, with a statistically significant increase in the levels between 28 and 32 h, as well as 32 and 36 h. A plateau in the BiP levels was observed between 40 and 44 h. Overall, all three proteins show a general increase over the 24-hour period examined.

#### Normalized protein levels

Given that the amount of all three proteins generally increases over the early trophozoite to mid schizont stage, it is plausible that the increase in PM I and PM IV expression may be accounted for by an increase in total protein per parasite as the trophozoite expands. Therefore, to account for total protein increase, the PM I and PM IV protein bands were normalized to BiP to show changes in the *relative* amounts of PM I and PM IV with respect to the total protein in the parasite (Figure 3D). In doing so, a steady expression relative to BiP was observed across timepoints, except for a small but statistically significant difference for PM I between 32 and 36 h. Therefore, within the limitations of the immunoblotting technique, the combination of the observed trends suggest that PM I and PM IV expression increases over the trophozoite to schizont stage in accordance with a universal cellular increase in protein levels as the parasite grows.

However, the time-dependent concentration of these enzymes in the DV is more insightful than absolute amounts with respect to the process of hemoglobin degradation within the DV. Therefore, the relative levels of PM I and PM IV were overlaid and compared to DV lumen growth (Extended Data Figure 4C-D) as well as the uptake into the DV (Extended Data Figure 4E-F). For both PM I and PM IV, the increase in relative protein abundance from 20 to 40 h corresponded with the increasing DV lumen volume and increase in uptake. The 44 h timepoint showed the largest difference between both PM I and PM IV levels and the DV lumen volume and uptake, respectively, compared to the other timepoints, given that the DV had already collapsed substantially. As a representation of PM concentrations, the levels of PM I and PM IV were then normalized to the DV lumen growth (Extended Data Figure 4G), where constant concentrations were observed between 28 and 40 h, but larger values obtained for the 20 to 28 h, and 44 h timepoints.

### Basal levels of the heme-containing species

#### Percentage and absolute levels

To probe the relationships between the DV lumen or proteases, and the basal levels of hemoglobin, heme and hemozoin, parasites were subjected to a time-dependent heme fractionation assay over a 24-hour period. The percentage levels for each heme-containing species in NF54 parasites remained stable over the entire 24-hour period resulting in an overall mean percent for hemoglobin (1.1 ± 0.1%), heme (7.9 ± 0.2%) and hemozoin (91 ± 0.2%) across all timepoints (Figure 4A-B). The absolute levels of hemoglobin, heme and hemozoin and the total Fe per parasite allow for direct comparisons between each heme-containing species, with those of hemoglobin Fe spanning a narrow range from 0.28 ± 0.02 fg/cell (20 h) to 1.3 ± 0.2 fg/cell (44 h). In contrast, the heme, hemozoin and total Fe levels appear to follow an exponential increase (Figure 4C-D), although the rate of this increase slows for both hemozoin and total Fe between 36 and 44 h, indicating more of a gradual sigmoidal profile. Given that hemozoin makes up approximately 90% of the total Fe, it is unsurprising that the trends for these two species resemble one another.

**Figure 4.**
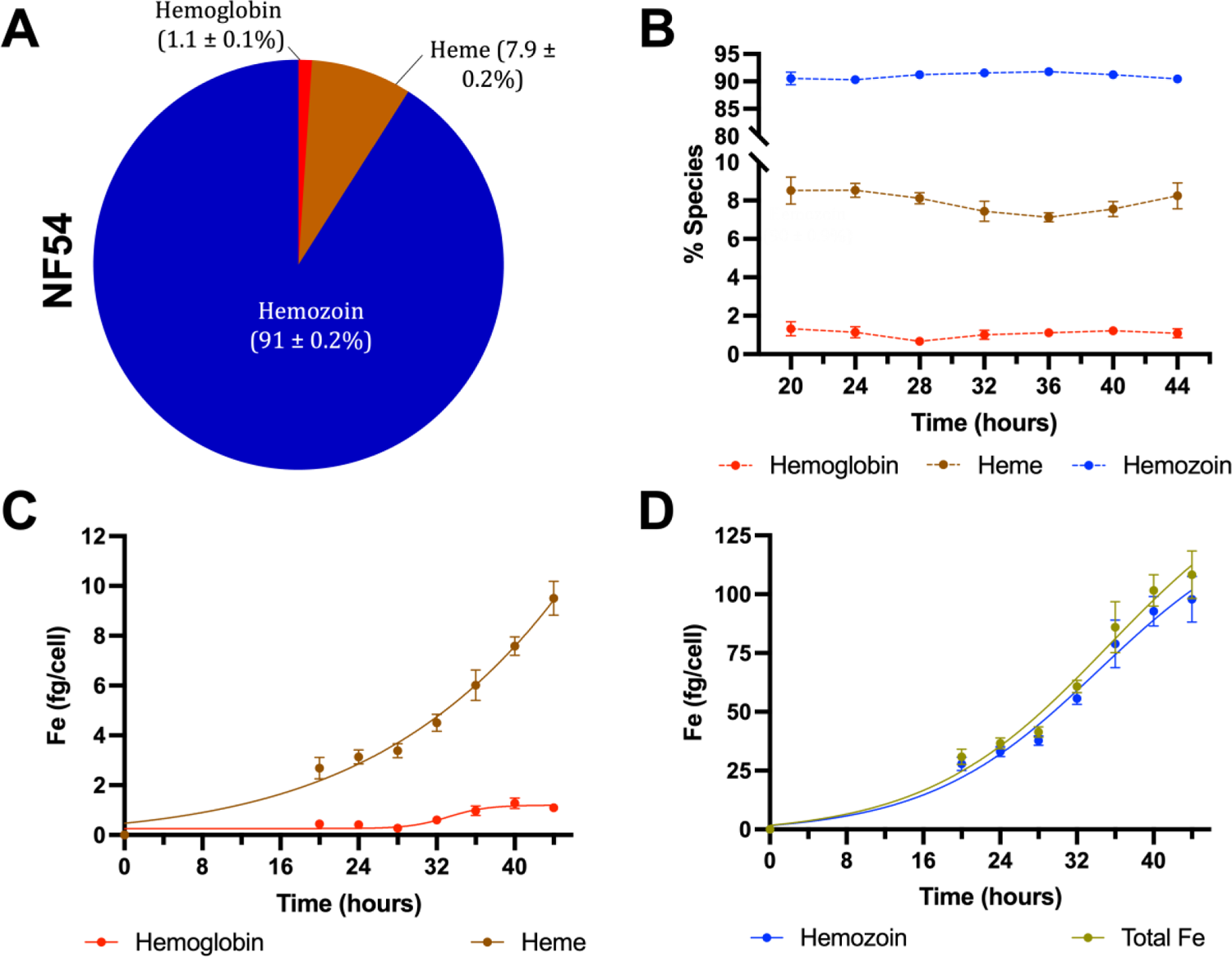
Time dependent NF54 cellular fractionation for describing the basal percent and absolute amounts of the heme-containing species over the trophozoite to schizont phases. **A**. The overall percent average of each heme-containing species across all timepoints and **B**. Percent heme-containing species over time. Using flow cytometry counts, the absolute amounts in fg/cell were calculated of **C**. hemoglobin and heme, **D**. hemozoin and total Fe.

Similar data was generated in the Dd2 strain and these showed comparable trends to NF54 with some minor differences (Extended Data Figure 5). First, the percent hemoglobin in Dd2 showed more variability and was larger than NF54 in the earlier trophozoite stage, resulting in a slightly larger overall percent of hemoglobin (3.5 ± 0.2% vs 1.1 ± 0.1%) and lower percent of heme (6.1 ± 0.2% vs 7.9 ± 0.2%) compared to NF54 (Extended Data Figure 5A-B). Interestingly, the maximum absolute amount of hemoglobin was also higher in Dd2 compared with NF54 (2.4 ± 0.4 vs 1.3 ± 0.2 fg/cell) and maximum absolute heme levels were lower (5.8 ± 0.9 vs 9.5 ± 0.6 fg/cell). Furthermore, the Dd2 curves for hemozoin and total iron increased exponentially across the entire 24-hour period, without reaching a plateau as was observed in NF54 (Extended Data Figure 5C-D).

### Heme detoxification insights

Given that the PMs and heme-containing species examined localize to the DV, additional calculations were made to investigate the relative concentrations (as opposed to absolute amounts) of the soluble species within the NF54 DV lumen, i.e. the PMs, hemoglobin and heme divided by the DV lumen volume measured herein (Figure 5A and Extended Data Figure 4G). The hemozoin, although within the DV, does not occupy lumen space and was therefore excluded from this analysis. Notably, during the NF54 mid trophozoite stage (28 to 40 h), the relative concentration of the soluble species remained constant across timepoints. On the other hand, the relative concentrations of these species were comparably higher during the early trophozoite phase (20 to 28 h) and the schizont stage (40 to 48 h). Similar calculations were applied to Dd2, and it was found that between 20 and 32 h the relative concentrations of hemoglobin and heme was lower compared to the period between 32 and 40 h.

**Figure 5.**
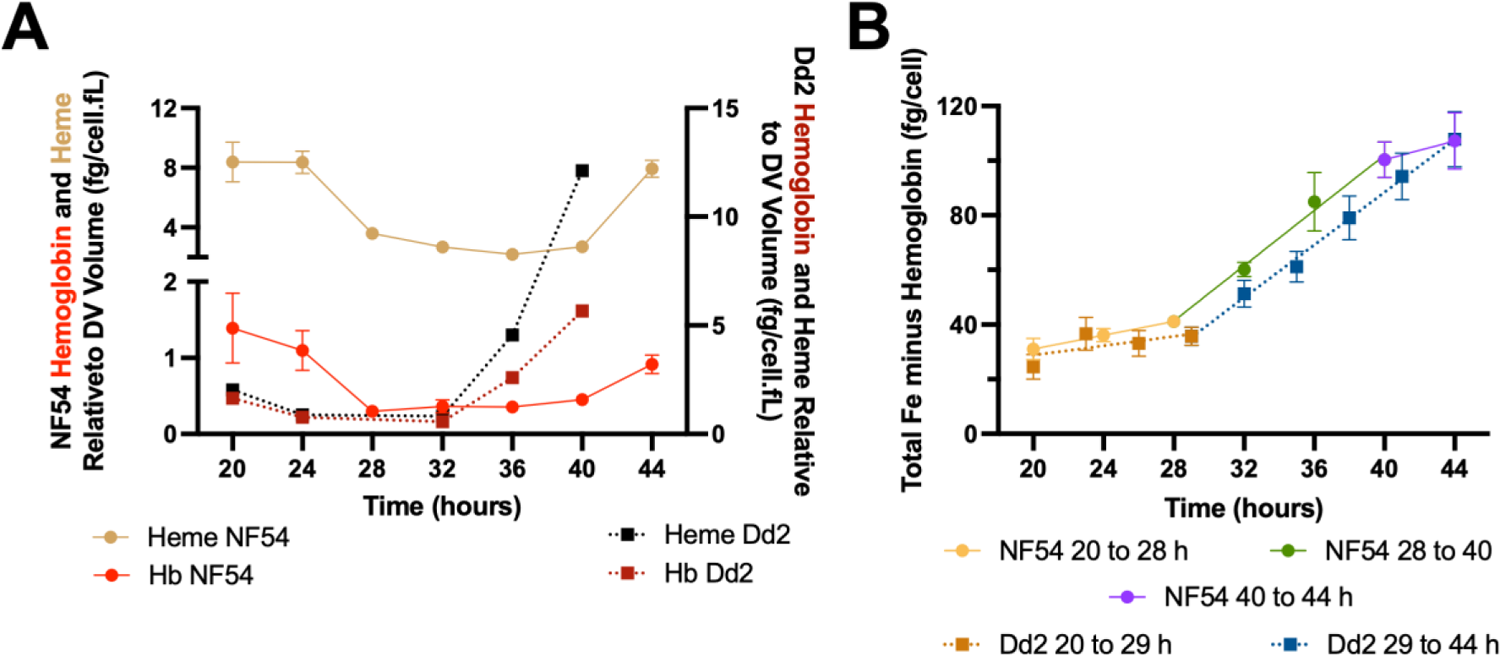
**A.** The amounts of hemoglobin and heme obtained from the heme fractionation experiments (Figure 4C and Extended Data Figure 5C), respectively, normalized to the DV lumen volume (Figure 2B) in both NF54 (circles, shown on left y-axis) and Dd2 (squares, shown on right y-axis). **B.** The amount of hemoglobin digested calculated as the Total Fe minus the absolute hemoglobin fraction in NF54 parasites obtained in the heme fractionation experiments (Figure 4C-D, and Extended Data Figure 5 C-D).

To further probe the degradation of hemoglobin, the amount of Total Fe minus the remaining hemoglobin fraction was calculated (Figure 5B). This metric corresponds to the amount of hemoglobin digested in femtograms (i.e. the heme plus hemozoin fractions). The gradient of the line therefore indicates the rate of hemoglobin digestion in femtograms per hour. Between 20 to 28 h, 28 to 40 h and 40 to 44 h, the rates were 1.3 fg/h, 5.1 fg/h and 1.7 fg/h, respectively, in NF54. Interestingly, in Dd2 parasites only two distinct phases were observed between 20 to 29 h and 29 to 44 h with digestion rates of 0.9 fg/h and 4.8 fg/h.

## Discussion

The broad aim of this work was to probe important DV-related features that affect hemoglobin catabolism in its native state in *P. falciparum* parasites. In turn, this provided a deeper understanding of the underlying parasite biology involved in the heme detoxification pathway. Four crucial pieces of data were analyzed across the trophozoite to schizont life stage: (i) the growth of the DV lumen volume, (ii) the uptake into the DV, (iii) the levels of PM I and PM IV proteases and (iv) the basal levels of hemoglobin, heme and hemozoin in untreated parasites.

It was found that the growth of the DV lumen best fit a Gompertz growth curve during the early-to-late trophozoite stage before collapsing during the schizont stage in both NF54 and Dd2 strains (Figure 2B). At the resolution of the confocal microscopy experiments used (Figure 2A), accurate volumetric analysis of the pre-DV acidic cytostomal invaginations^14–18^ was not feasible. Thus, data was not collected between 0 and 20 hours, and the assumption was made that the growth would follow the fitted curve during this period starting at (0,0). The uptake of pHrodo™ into the DV followed a comparable trend to the lumen volume, with the total uptake increasing dramatically from the early trophozoites to the late trophozoite stage, and then decreasing within the schizont stage (Figures 2C), while the concentration of pHrodo remained constant (Figure 2D). This indicates that lumen growth and uptake are tightly correlated, such that as the lumen grows, the parasite takes up more material leading to a constant pHrodo™ concentration within the vacuole.

This sigmoidal-type growth of the lumen and uptake in both strains can be rationalized by considering the basic underlying biology of the parasite postulated by Tilley and colleagues that during the late ring stage, acidic, endocytic pre-DV vacuoles fuse to initiate DV biogenesis.^16^ If one were to approximate the DV by a sphere, the volume would be calculated as: V = ^4^ πr^3^, while the surface area would be calculated as: SA = 4πr^2^, where r is the radius of the sphere. This indicates that the volume, which is dependent on r^3^, would grow faster than the r^2^-dependent surface area. In the context of the DV, the surface area would limit how quickly the volume grows because the surface of the DV provides the contact point for fusion of new vesicles. The larger the surface area, the more vesicles can fuse and therefore the larger the DV can grow. Between 0 and 20 h, the parasite is present in the ring phase, and therefore, the pre-exponential phase seen towards this asymptote is likely attributed to the limited surface area available as the small individual vesicles combine to form the central DV compartment seen in the trophozoite stage. This period (0 to 20 h) also corresponds to a lag in uptake which reflects the low metabolic energy demand of the ring stage.

The growth phase between 20 and 36 h, and 20 and 32 h in the NF54 and Dd2 strains, respectively, corresponds to the trophozoite stage of the parasite. During this stage, the numerous small pre-DV vesicles continue fusing with the central DV compartment. The addition of each small vesicle in 3D space results in an ever-increasing surface area and therefore rapid volume growth, giving rise to the exponential-phase Gompertz growth and uptake curves. Notably, this lumen growth phase after 20 h corresponds to the most metabolically active stage of the parasite,^55^ correlating with the need for the rapid increase in uptake of RBC cytosol and large increase in lumen volume. As the energy demand of the parasite decreases and the majority of the hemoglobin from the RBC cytosol has been digested, the uptake and the growth begin to slow down resulting in the curve reaching its upper asymptote. This asymptote is visible in Dd2 for both the growth and uptake, but NF54 does not appear to reach an asymptote before the DV lumen begins to collapse. One possible explanation for this difference is that the Dd2 DV grows faster than that of NF54, thus reaching a point where the volume to expand is limited sooner than in NF54. It therefore approaches the upper asymptote more gradually as space within the RBC becomes limited.

As the parasites enter the schizont stage at approximately 36 to 40 h in both strains, the DV lumen begins to collapse. This does not preclude the existence of the DV, rather the data suggests that during this schizont phase, the DV mainly consists of hemozoin crystals and very little intermembrane space (compare T36 and T44, Figure 2A, and T36 and T40, Extended Data Figure 3A). Further evidence for the final collapse of the lumen can be seen in the schizont TEM image (Figure 2B, Schizont **Inset**), which contains merozoites and shows the hemozoin crystals tightly wrapped in the DV membrane (white arrow). This residual membrane-wrapped hemozoin is left behind when the schizonts burst to release merozoites and can be observed as debris under a light microscope (not shown).

A previous study has shown that the total DV volume, which includes hemozoin crystals, of the wild-type HB3 and drug-resistant Dd2 strains follows a sigmoidal growth pattern.^44^ While the shape of the Dd2 growth curve herein agrees with the previous study, the observed absolute volumes differ.^44^ The maximum total volume of the Dd2 DV, including hemozoin crystals, was previously shown to reach ∼5.5 fL in the late-trophozoite stage (30 h), while the volume for the Dd2 DV lumen in our study was shown to reach 3.7 fL at a similar stage (32 h), a difference of 1.8 fL. This variance is likely attributed to the volume occupied by the hemozoin crystals, which are excluded in the current measurements and appear from this analysis to occupy almost a third of the DV at this time point. Furthermore, the previous study did not present data beyond 30 h and therefore the collapse of the DV lumen was not studied. This collapse may be attributed to several factors. In light of the fact that the growth of the total DV volume as well as the hemozoin crystals themselves reach a plateau in the late trophozoite,^44, 56^ if the growth kinetics of the hemozoin crystals overtake the growth kinetics of the DV, this would lead to the hemozoin crystals taking over the available lumen space within the DV, resulting in the observed lumen collapse. A second possibility is that as the RBC cytosol uptake decreases in the schizont stage, the volume occupied by hemoglobin decreases which then manifests as a collapsing DV. An additional unknown is the fate of the contents of the DV, which are possibly exported into the cytoplasm, allowing vital components to compartmentalize into the forming merozoites.

Interestingly, it was observed that the volume of the lumen in Dd2 was larger than in NF54 across all timepoints, similar to previous observations made between the total DV volumes of drug-sensitive and resistant strains.^44, 57–59^ These studies attributed the observed difference to *Pf*CRT mutations. The association of *Pf*CRT mutations with altered vacuolar volume was previously confirmed using a CQS strain transfected with mutant *Pf*CRT, C3^Dd2^.^44^ This isogenic edited *Pf*CRT mutant showed increases in DV volume comparable to Dd2 relative to an unmodified CQ sensitive laboratory strain, GCO3, implicating the direct or indirect involvement of *Pf*CRT.^44^ Recent observations that the native function of *Pf*CRT involves transport of key substrates (such as peptides), which appears to be impaired in mutant *Pf*CRT, supports its role in causing larger vacuoles in drug-resistant parasites relative to drug-sensitive parasites.^60–62^ Indeed, this study suggests that the increased total DV volume observed in *Pf*CRT mutants is as a result of increased lumen volume and not merely more hemozoin. Thus, the impaired function of mutant *Pf*CRT may cause peptides that would otherwise be exported to accumulate within the DV lumen of Dd2 parasites resulting in the increased lumen. Additionally, we observed that Dd2 parasites have a higher basal level of hemoglobin than NF54 (Figures 4A-C and Extended Data Figures 5A-C), suggesting that the Dd2 lumen contains more hemoglobin at any given time compared to NF54. Therefore, one hypothesis is that the inhibited transport of substrates by mutant *Pf*CRT in Dd2 not only leads to larger vacuoles, but causes a shift in the equilibrium between the digested vs undigested hemoglobin, relative to NF54.^44, 63^

For further insights and a holistic view of the heme detoxification process, the inter-strain levels of all the heme-containing species within the DV were investigated. When looking at the overall percentage of each species over the entire stage (Figure 4A and Extended Data Figure 5A), it appears that in general Dd2 parasites not only had a higher percentage of hemoglobin, but also a lower percentage of heme compared to NF54 parasites, while the percentage of hemozoin was comparable between the strains. Additionally, it was interesting to note that although heme has been shown to be toxic to the parasite,^64, 65^ it can tolerate seemingly non-toxic levels as the DV lumen volume expands, since the total heme concentration remains below 13 fg of heme Fe per fL (Figure 5A) or 0.233 M.

While the percentage of each heme-containing species remained steady over the trophozoite to schizont life stage, a time-dependent increase in the absolute values of all species was observed in both NF54 and Dd2 strains to varying degrees (Figure 4 and Extended Data Figure 5). Prior to 20 h, ring stage parasites cannot adequately pellet during centrifugation to give heme-species data points and therefore the absolute level graphs were interpolated with the point (0,0) and extrapolated using this point between 0 and 20 h. The biological basis of the interpolation of this point is that merozoites have been shown to contain an inactive cytostomal ring on their periphery, indicating that parasite feeding has not yet commenced at this stage.^14, 66^ This extrapolation results in a gradual increase in heme-containing species defined by the curve fit between 0 and 20 h, corresponding to the ring stage. The extrapolation of the hemoglobin fraction is supported by the presence of active cytostomes during the ring stage, allowing the parasite to begin feeding on the RBC cytosol pre-trophozoite phase^67^ Support for the extrapolation of the heme and hemozoin curves, lies in the fact that previous studies have documented the presence of pre-DV, acidic vacuoles that begin to form during the ring stage which have been shown to contain hemozoin, substantiating the catabolism of hemoglobin in the early ring stage.^16, 68, 69^ Naturally, these factors would in turn support the extrapolation of the total Fe curve.

The absolute hemoglobin levels remained within a narrow range in both NF54 and Dd2, while the heme, hemozoin and total Fe levels showed exponential increases to varying degrees (Figure 4C-D and Extended Data Figure 5C-D). A more complete picture of the pathway emerges when the percent and absolute amounts are considered holistically and in combination with the DV lumen volume (Figure 5A). In general, there is an interdependence between the growth of the DV lumen volume and the absolute amounts of the heme-containing species such that the proportion of each species (Figure 4B and Extended Data **5B**) remains approximately constant. This, coupled with the fact that the heme concentration remains below 233 mM, indicates a tightly regulated homeostasis of these species that the parasite maintains throughout the trophozoite to schizont life stage.

Notably, in NF54, the total Fe over the 24-hour period (Figure 4D) and the uptake of pHrodo into the DV (Figure 2C) corroborate each other. In NF54, from 20 to 40 h the amount of RBC cytosol taken up and total Fe increases exponentially, after which the DV collapses, less cytosol is taken in and the rate of increase in the total Fe begins to level off (Figure 2C and **4D**). This is unsurprising given that the source of the heme-derived Fe in the parasite is derived predominantly from hemoglobin uptake and digestion.^70^ In Dd2, however, although uptake into the DV plateaus before decreasing (Figure 2C), the total Fe increases exponentially over the same period (Extended Data Figure 5D), further supporting the disruption caused by *Pf*CRT mutations to processes in the DV, relative to the wild-type. A noteworthy observation is that the low levels of hemoglobin and gradual increase in total Fe over time do not support one of the modes of uptake previously suggested termed the “big gulp”.^18^ Rather the data support the reliance on cytostomal invaginations as the primary method of uptake in line with other previous findings that have also contradicted the big gulp theory.^14, 16–18, 69, 71, 72^

An interesting basis for how hemoglobin is processed in wild-type parasites is provided by a holistic examination of the data. Over the trophozoite to schizont stage (28 to 40 h) a tight homeostasis of hemoglobin degradation and the heme detoxification process is maintained in NF54 parasites (Figure 5A). On either side of this stage, early trophozoite (20 to 28 h) and early-to-late schizont (40 to 48 h), there appears to be higher relative concentration of the PMs (Extended Figure 4G), hemoglobin and heme (Figure 5A), however, there also appears to be a slower rate of hemoglobin catabolism compared to the mid-trophozoite stage (Figure 5B). During these periods the DV lumen is at its smallest and correlates with stages when the parasite and therefore, the DV, are transitioning between different phases, i.e. from ring to trophozoite in the early stage and from trophozoite to schizont in the latter. During the mid trophozoite stage (28 to 40 h), the relative concentration of the soluble material appears to be lower, but constant, while the hemoglobin degradation rate appears to be higher. This correlates with the period in which the parasite is most metabolically active and reliant on hemoglobin degradation and heme detoxification. Most interestingly, is that regardless of the phase, the proportion of the hemoglobin, heme and hemozoin relative to each other remains constant.

In Dd2, a slightly different pattern is observed. It appears that rather than three distinct phases, there are two. The earlier-to-mid trophozoite stage (20 to 32 h) correlates with lower relative concentrations of hemoglobin and heme (Figure 5A), as well as a lower rate of hemoglobin degradation (20 to 29 h, Figure 5B). The relative hemoglobin and heme concentrations then increase past 32 h correlating with a higher rate of hemoglobin degradation. Despite these observed differences, similar to NF54, the proportions of the heme-containing species remain constant.

When considering all the parameters studied, these data contribute to a holistic overview of the heme detoxification pathway which provides a deeper understanding of the fundamental biology of the DV. A theoretical DV physiology schematic for NF54 and Dd2 are shown in Figure 6 and provide insights into wild-type and CQ-resistant parasites. Included in this figure is a summary of the data presented in this manuscript. The picture of the NF54 DV that emerges is one that is characterized by three distinct growth periods: (1) lag-type growth (20 to 28 h), (2) rapid growth (28 to 40 h), and (3) the plateau (40 to 48 h).

**Figure 6.**
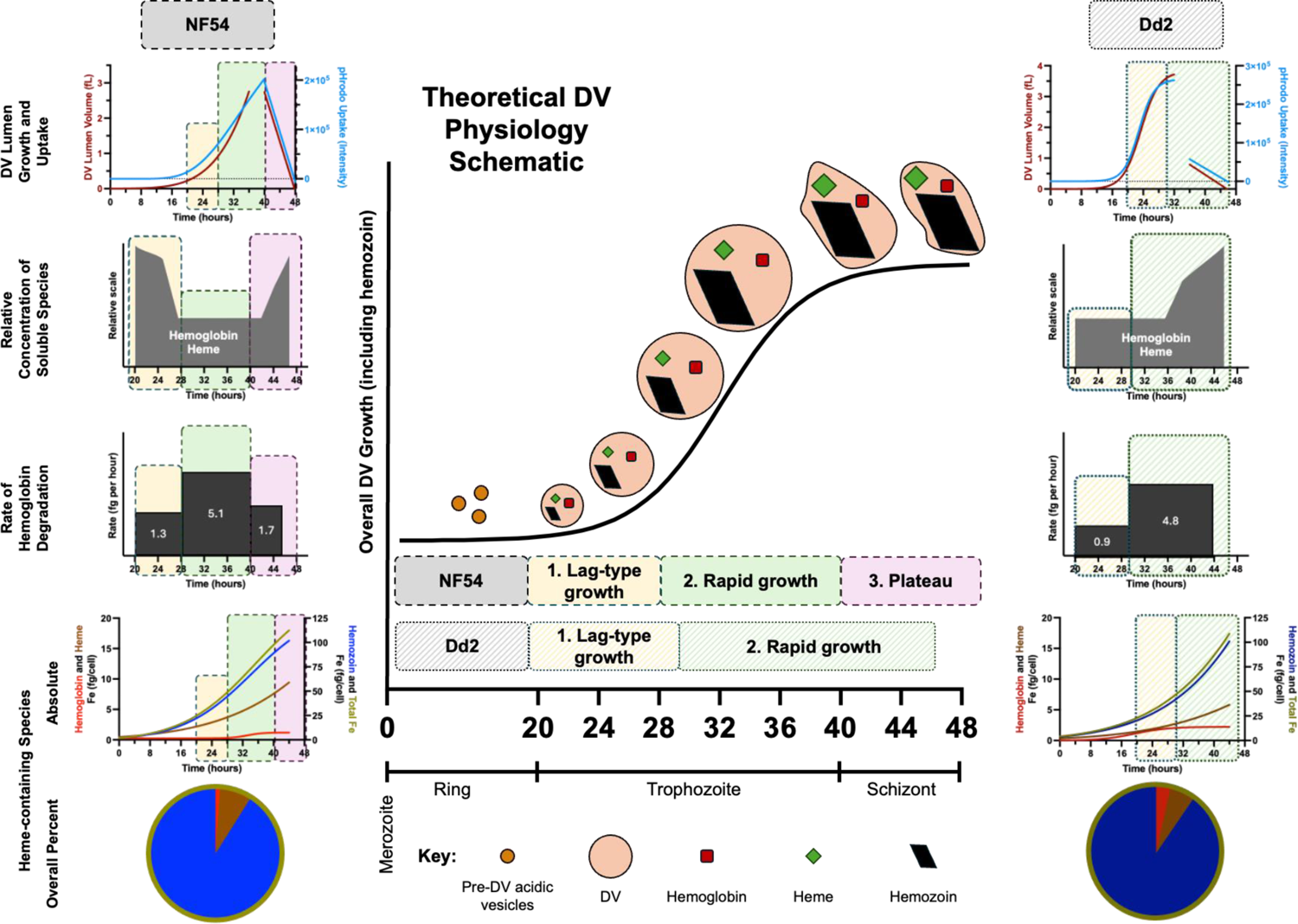
Proposed DV physiology and summary of the experimental input data required for the modelling of the heme detoxification pathway showing the growth phases for NF54 and Dd2 parasites.

During the lag-type growth (20 to 28 h) the lumen volume, uptake into the DV and levels of heme, hemozoin and total Fe increase gradually. An interesting feature of this growth period is the relatively higher concentration of the soluble matter within the DV compared to later stages. It is unclear why the concentration level of the soluble matter would be elevated compared to the later stages; however, this may be due to a concentrating of the soluble matter within a relatively smaller DV and a hallmark of a transition period of the parasite DV. The rate of hemoglobin degradation is lowest during this phase.

The lag-type growth period is followed by the rapid growth period (28 to 40 h) which is characterized by a significant increase in the DV lumen volume, uptake and levels of each heme-containing species. This corresponds to the most metabolically active stage of the intraerythrocytic parasite life cycle with the largest contribution to overall ABS parasite growth, where increased hemoglobin catabolism allows for the preservation of intracellular osmotic pressure. ^37, 38^ During this mid-trophozoite phase there are higher relative protein levels of PM I and PM IV compared to the earlier phase, which would allow for the processing of increasing amounts of hemoglobin, however, a steady state concentration of these proteins maintains the constant concentration of hemoglobin, characterized by a higher rate of hemoglobin degradation compared to the earlier or later stages.

The final phase, the plateau (40 to 48 h), is distinguished by a collapse of the DV lumen and a linear decrease of the uptake into the DV. The decreased uptake is reflected in the slowing increase of the levels of hemozoin and total Fe in NF54. The heme levels on the other hand continue to follow the exponential curve trajectory, presumably as a result of continued hemoglobin degradation. At this point in the life cycle, the parasite has grown to a size that takes up most of the space within the RBC and therefore, its dependence on hemoglobin uptake becomes inconsequential, and parasite resources are diverted towards schizogony and merozoite formation leading to a lower rate of hemoglobin degradation. The penultimate fate of the DV is a residual body of “shrink-wrapped” hemozoin crystals.

In contrast, although the growth of and uptake into the Dd2 DV mirrors the NF54 DV, the underlying hemoglobin degradation characteristics indicate two distinct phases: (1) lag-type growth (20 to 29 h) and (2) rapid growth phases (29 to 46 h). During the first phase, the relative concentration of soluble matter remains lower compared to the latter phase, while the increase in the heme-containing species is gradual. During the second phase, the heme-containing species, except hemoglobin, increase dramatically accompanied by large increases in the relative concentration of the soluble species. Finally, the rate of hemoglobin degradation appears to be lower during the first phase and higher during the second. Overall, these differing observations support the observations that *Pf*CRT mutations in Dd2 parasites affect hemoglobin processing relative to wild-type parasites.

This study presents a holistic overview of the physiology of the malaria parasite DV, an important organelle and site of action of numerous clinical and experimental antimalarial drugs. Given that important ACT partner drugs and those in development target hemozoin formation, understanding the mechanics of this pathway remain imperative. This study also highlights some of the strain differences between wild-type NF54 and CQ-resistant Dd2 strains in terms of DV lumen volume, uptake into the DV and basal levels of heme species over time. Thus, the continued study of the heme detoxification pathway and DV physiology may lead to a basis for an improved understanding of resistance mechanisms. Indeed, these acquired data provide a platform and framework to extend the current knowledge of DV physiology and thus, beckons the continued study of the vital heme detoxification pathway at a basal level between different drug-resistant parasite strains. Finally, understanding how pathway inhibitors affect hemoglobin processing is equally important and would allow for specific parts of the pathway to be further probed.

## Author Contributions

L.F.G.; conceptualization, investigation, analysis, writing original draft, reviewing and editing. T.J.E.; funding acquisition, conceptualization, supervision. K.J.W.; funding, supervision, writing, reviewing and editing.

## Acknowledgements

This research is dedicated to the late Professor Timothy Egan. The study was funded by the National Institute of Allergy and Infectious Diseases of the National Institutes of Health under Award Number 5R01AI143521. The content of this publication is solely the responsibility of the authors and does not necessarily represent the official views of the National Institutes of Health. We thank Dr. David Fidock and the Fidock Lab from Columbia University Irving Medical Center for donating the Dd2 knockout lines and plasmepsin antibodies. We would like to also thank Dr. Thomas Oelgeschläger and Dr. Jasmin Knopp from the Faculty of Molecular and Cell Biology at the University of Cape Town for the contribution of their expertise and training in western blot techniques, as well as the access to their laboratory and resources to conduct the research. We thank Dr Joanne Egan for the use of her TEM images in Figure 2B.

## Notes

### Competing Interest Statement

The authors have declared no competing interest.

