## Extended Data Figures and Methods for "Heme Detoxification in the Malaria Parasite *Plasmodium falciparum*: A Time-Dependent Basal-Level Analysis"

TABLE OF CONTENTS

#### Extended Data Figures

**Extended Data Figure 1.** Blood stage culture clock and flow cytometry standard of NF54 *P. falciparum* parasites.
**Extended Data Figure 2.** Blood stage culture clock and flow cytometry standard of Dd2 *P. falciparum* parasites.
**Extended Data Figure 3.** Digestive vacuole lumen volumetric and uptake analysis in Dd2 *P. falciparum* using confocal microscopy.
**Extended Data Figure 4**. Immunoblot analysis of trophozoite whole cell extracts.
**Extended Data Figure 5**. Time dependent cellular fractionation assays in the Dd2 strain.

#### Methods

**Culture Care** ........ 7
- General method
- Preparation of cultures for time-sensitive assays
- Percoll® gradient enrichment
- Flow cytometry

**Digestive vacuole studies** ........ 8
- RBC loading
- Incubation and harvesting
- Confocal imaging
- Image analysis

**PM I and IV protein analysis** ........ 9
- Parasite lysate preparation
- SDS-page
- Western blot preparation
- Densitometric blot analysis

**Basal levels of the heme-containing species** ........ 11
- Incubation and harvesting
- Cellular fractionation and heme standard curve

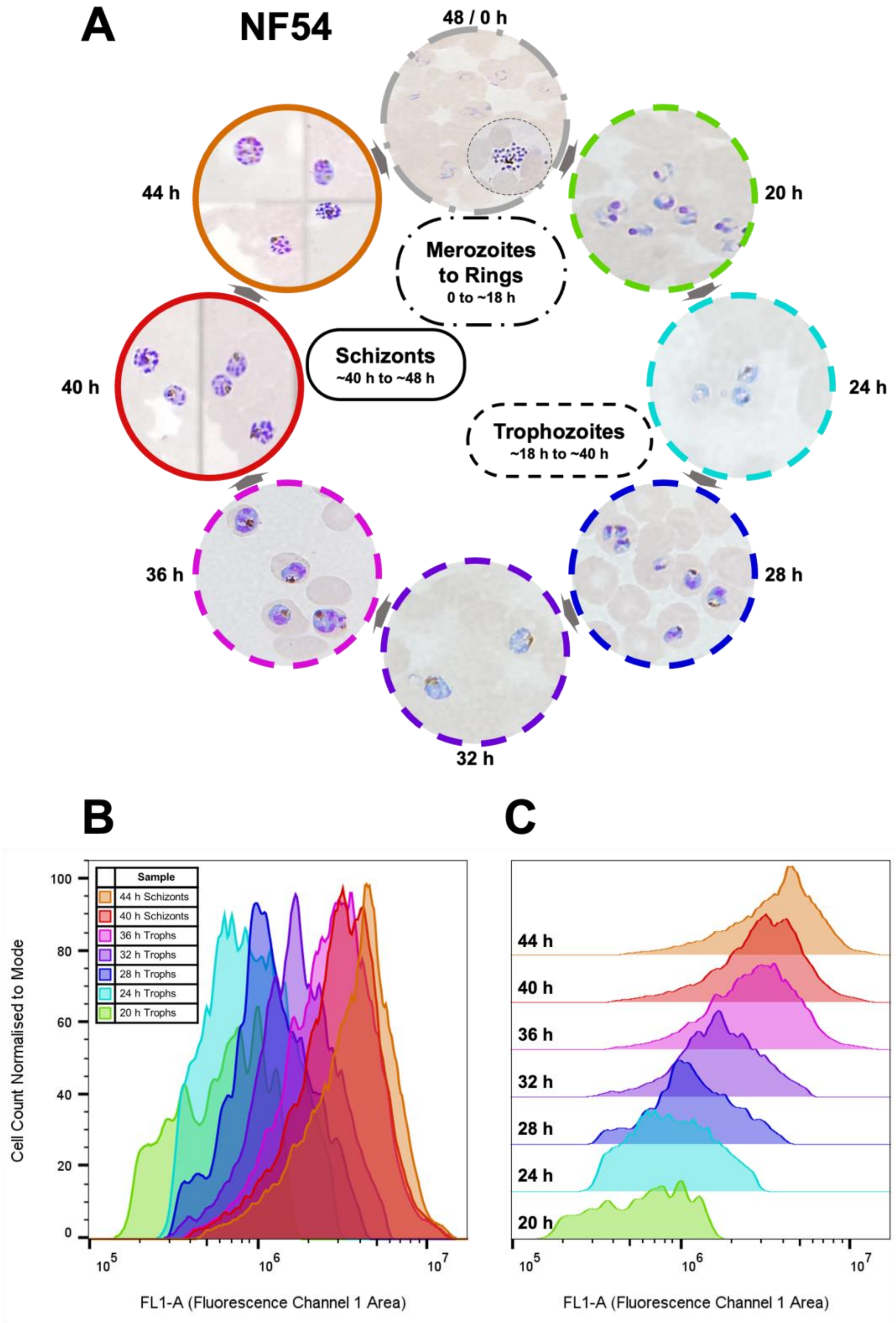


**Extended Data Figure 1.** Blood stage culture clock and flow cytometry standard of NF54 *P. falciparum* parasites. **A.** Giemsa-stained NF54 parasites shown in different phases as viewed under a light microscope. The ring and merozoite images were collected during routine culturing, while the 20 to 44 h images were collected at consecutive 4-hour intervals. **B.** Overlaid flow cytometry histograms of NF54 trophozoites stained with SYBR® Green collected at consecutive 4-hour intervals. **C.** Staggered histogram of Graph B**.** Experimental samples were compared visually to these standards to confirm parasite age.

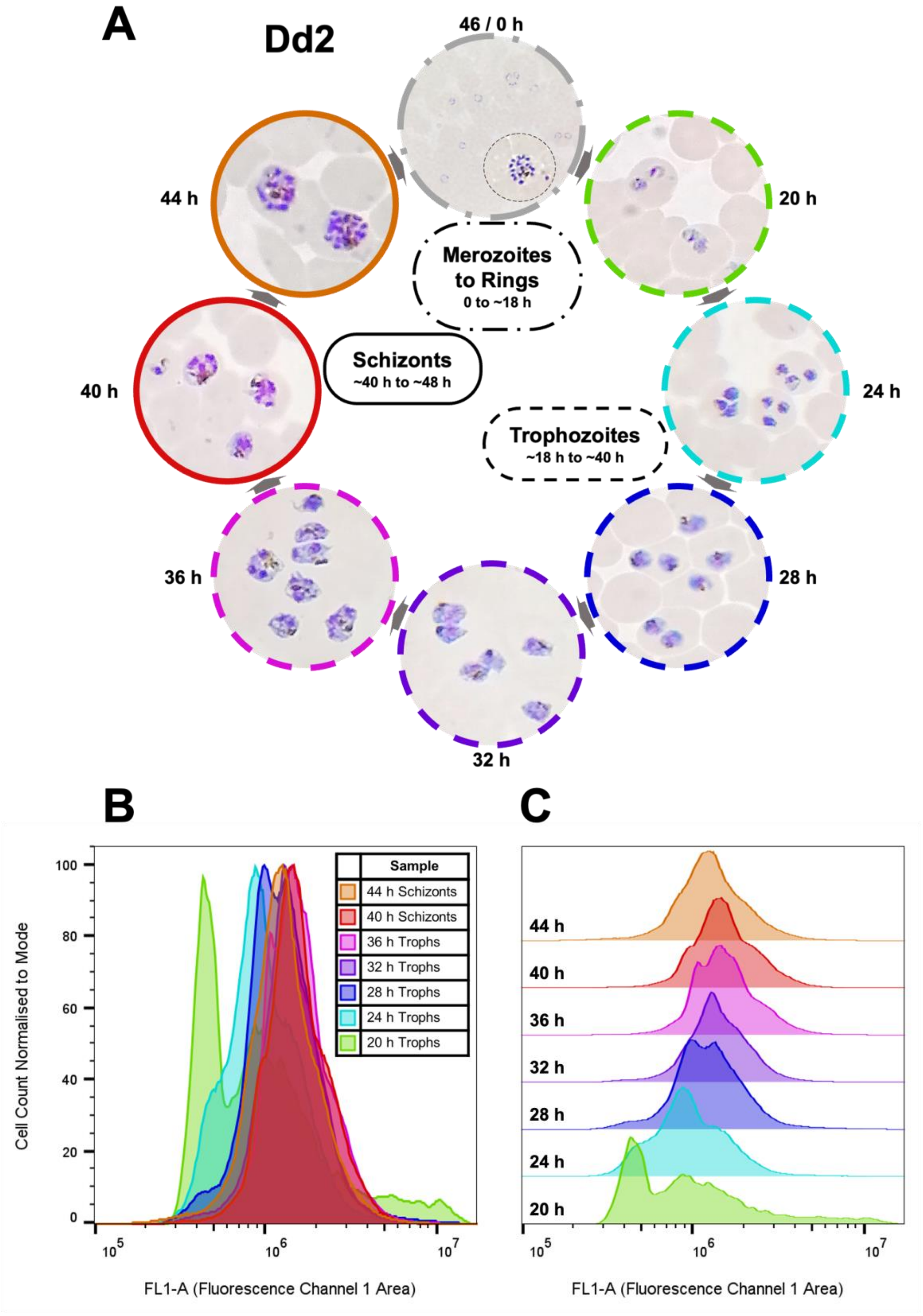


**Extended Data Figure 2.** Blood stage culture clock and flow cytometry standard of Dd2 *P. falciparum* parasites. **A.** Giemsa-stained Dd2 parasites shown in different phases as viewed under a light microscope. The ring and merozoite images were collected during routine culturing, while the 20 to 44 h images were collected at consecutive 4-hour intervals. **B.** Overlaid flow cytometry histograms of Dd2 trophozoites stained with SYBR® Green collected at consecutive 4-hour intervals. **C.** Staggered histogram of Graph B. Experimental samples were compared visually to these standards to confirm parasite age.

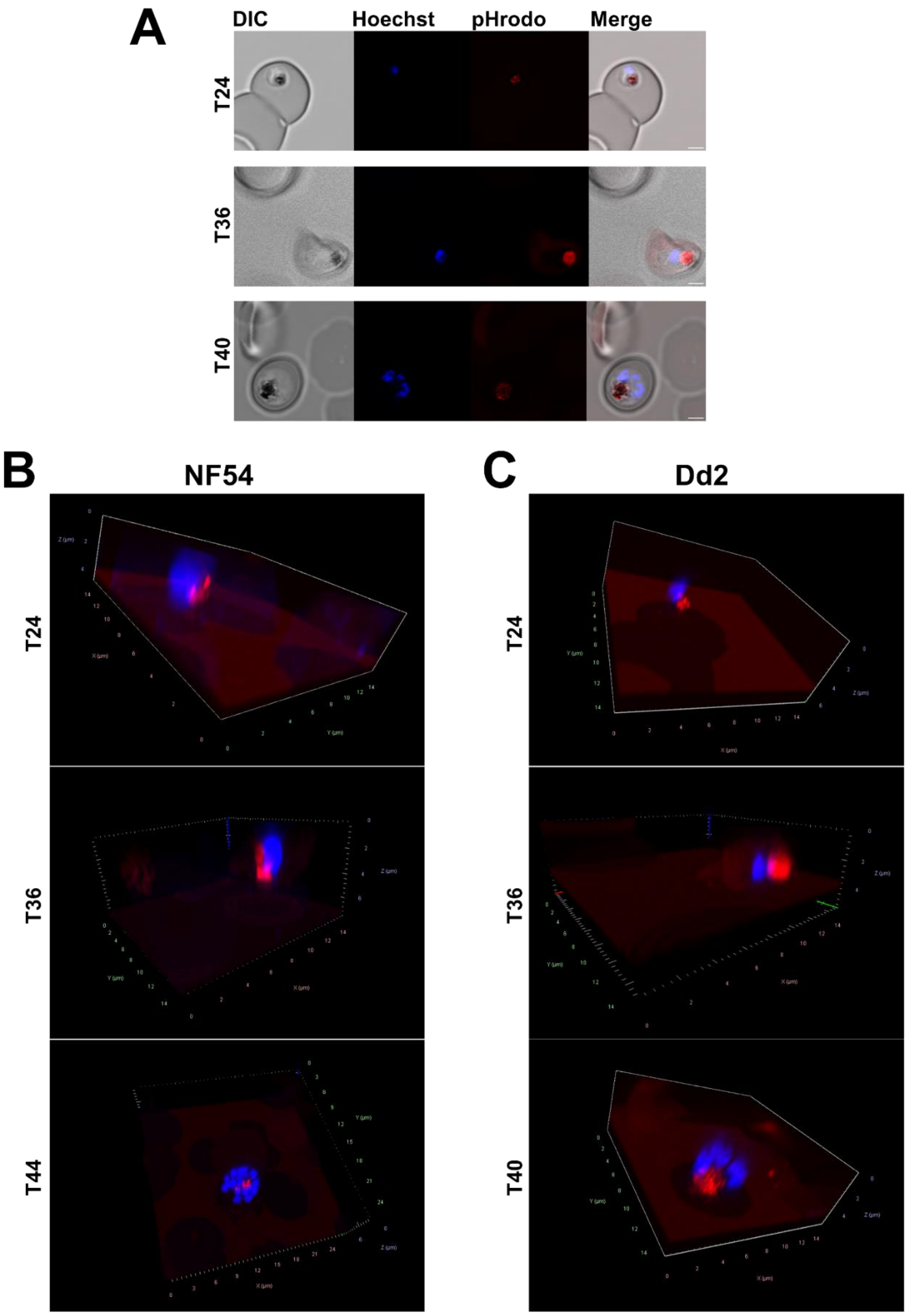


**Extended Data Figure 3.** Digestive vacuole lumen volumetric and uptake analysis in Dd2 *P. falciparum* using confocal microscopy. Parasites were allowed to re-invade RBCs pre-loaded with pHrodo™-dextran red and imaged using the Z-stack functionality of an LSM 880 Airyscan confocal microscope (ZEISS). **A.** A single Z-stack slice of representative individual RBCs infected with Dd2 *P. falciparum* parasites at various timepoints (stipulated on image). Hoechst (blue) stains the DNA while pHrodo™ (red) selectively stains the DV. All scale bars represent 2 µm. 3D renderings produced using Zen Blue (ZEISS) of the combined Z-stack slices of the individual **B.** NF54 (ED Figure **1A**) and **C.** Dd2 (shown in **A**) parasites at various timepoints.

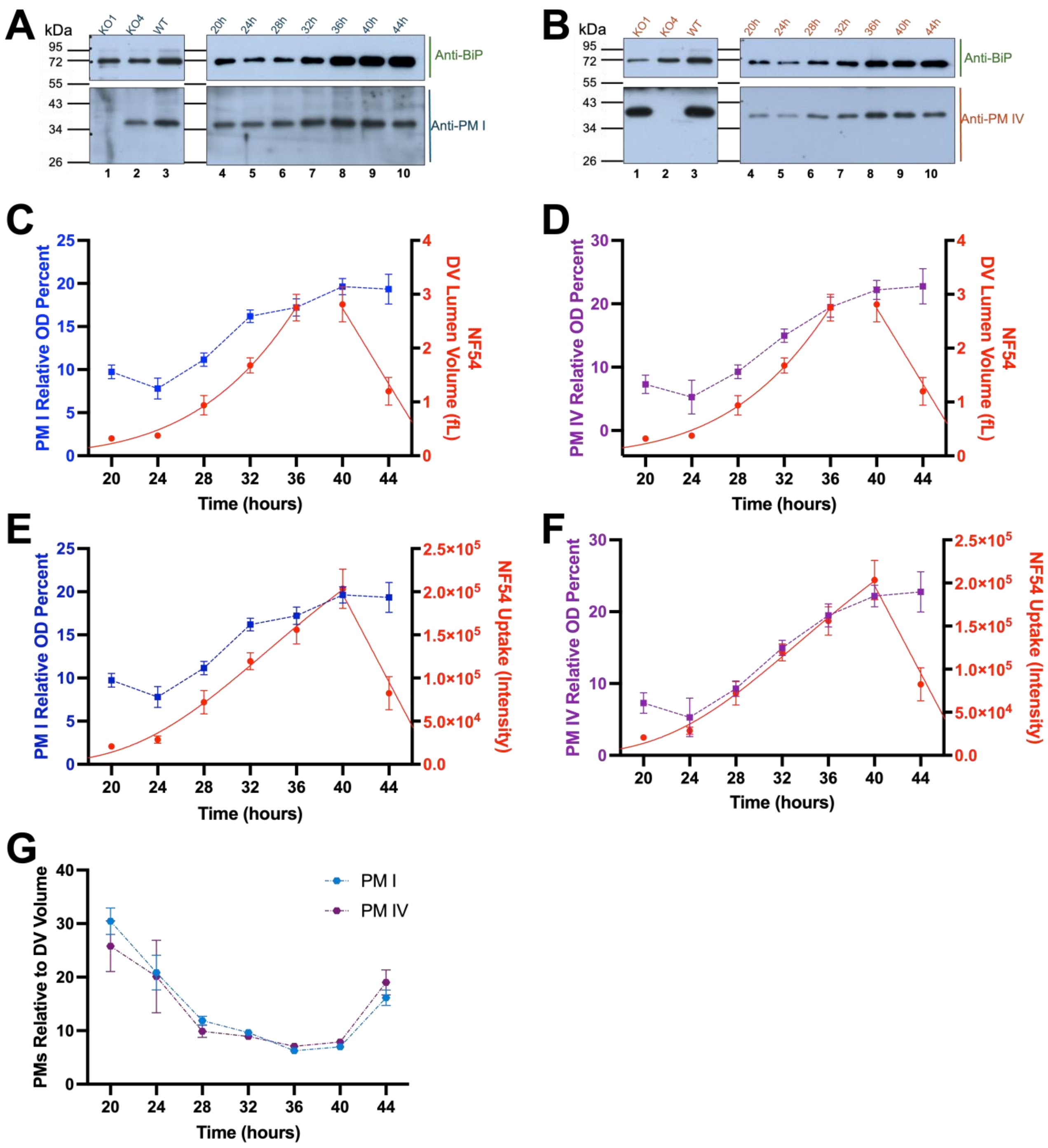


**Extended Data Figure 4.** Immunoblot analysis of trophozoite whole cell extracts from wild type NF54 (50 000 parasites per lane) harvested at 4-hour intervals along the trophozoite cycle (stored in 1× SAB), and Dd2-derived PM I and PM IV knockout lines, using anti-PM and anti-BiP antibodies. Proteins were separated by 12% SDS-page, transferred to a PVDF membrane, probed with anti-PM I (1:10 000), anti-PM IV (1:10 000) and anti-BiP (1:20 000), as indicated, and visualised by chemiluminescence on X-ray film. **A.** Immunoblot probed with anti-BiP (top panel) and anti-PM I (bottom panel). **B.** Immunoblot probed with anti-BiP (top panel) and anti-PM IV (bottom panel). Exposure times were optimised for each blot. The separated panels indicate where the membrane was separated for incubation with the primary antibodies. "WT" (lane 3) refers to the NF54 mixed trophozoite sample, "KO1" (lane 1) and "KO4" (lane 2) refers to the PM I and PM IV KO lines, respectively. The overlay of the relative protein abundance of **C.** PM I and **D.** PM IV in arbitrary units (left y-axis), determined by densitometric analysis as shown in Figures **3A-B**, with the DV lumen volume (right y-axis) shown in Figure **2B** over the trophozoite to schizont stage. The overlay of the relative protein abundance of **E.** PM I and **F.** PM IV in arbitrary units (left y-axis) with the uptake into the DV (fluorescence intensity on the right y-axis) shown in Figure **2C** over the trophozoite to schizont stage. **G.** The amounts of PM I and PM IV obtained from the immunoblot experiments (Figure **3 A-B**) normalized to the DV lumen volume (Figure **2B**). For each data point of the protein data N = 4, for each data point of the uptake n ≥ 15 (error bars indicate SEM).

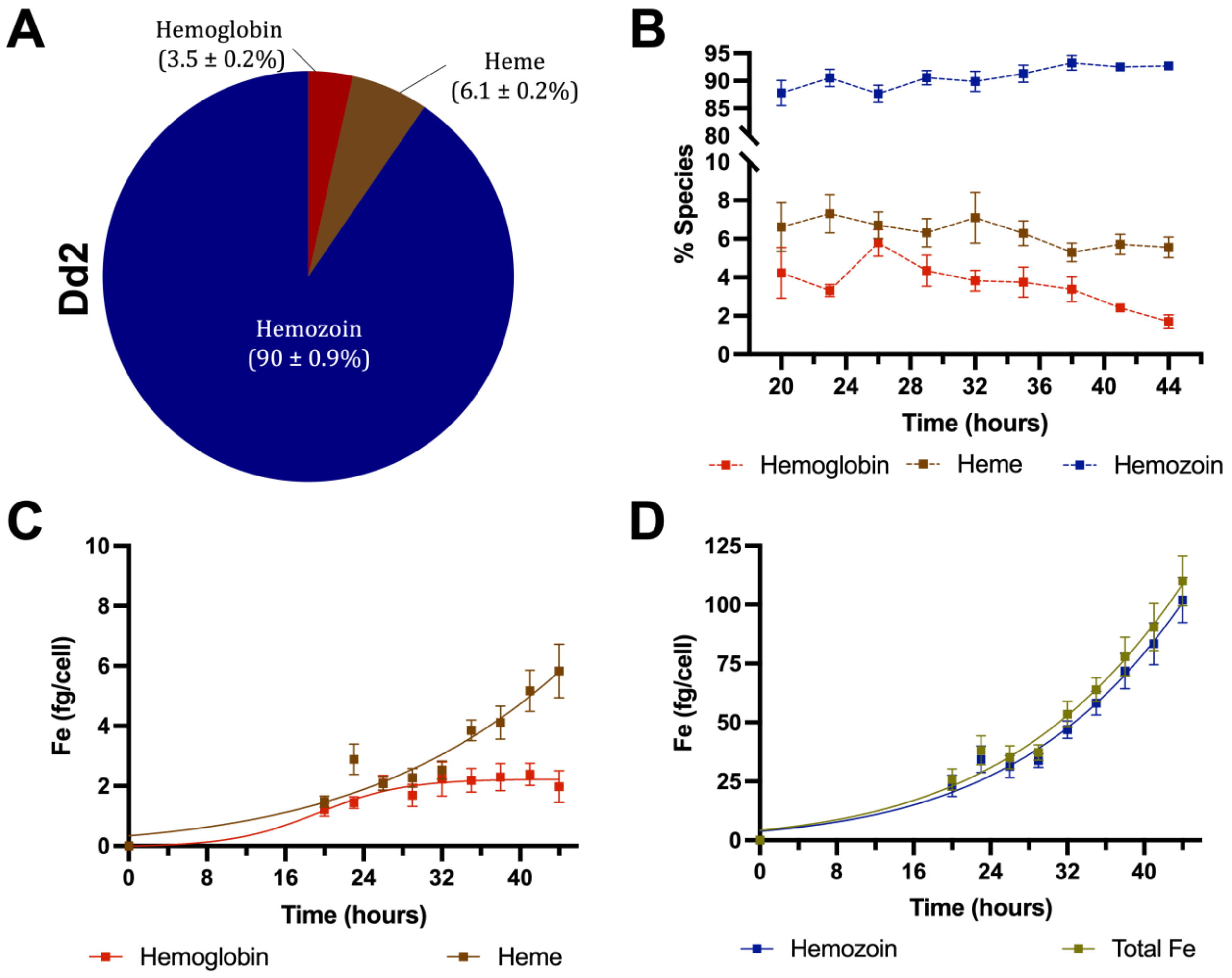


**Extended Data Figure 5**. Time dependent cellular fractionation assays were run using the Dd2 strain to determine the percent and absolute amounts of the heme-containing species over the trophozoite to schizont phases. **A**. The overall average of each species calculated as the average of the percent of the heme-containing species across all timepoints. **B**. Percent heme-containing species over time. Using flow cytometry counts, the absolute amounts in fg/cell were calculated of **C**. hemoglobin and heme, **D**. hemozoin and total Fe.

### Methods

*Cell culture*

**General method.** *Plasmodium falciparum* parasites were maintained in continuous culture based on the Trager and Jensen method.[1] Briefly, parasites were maintained in complete medium (10.4 g/L RPMI-1640, 4.0 g/L glucose, 0.088 g/L hypoxanthine, 6.0 g/L HEPES, 5.0 g/L Albumax II, 0.05 g/L Gentamycin and 2.25 g/L sodium bicarbonate) and $O^+$ red blood cells obtained from the Western Province Blood Services, Cape Town, at 5% parasitemia and 2% hematocrit. Ring stage parasites were synchronized by treatment with 5% sorbitol.[2]

**Preparation of cultures for time-sensitive assays.** In preparation for time-sensitive assays, parasite cultures were monitored closely to track parasite and kept tightly synchronized for at least one week prior to experimental set-up. For NF54 parasites, cultures were synchronized within two ranges in consecutive 48-hour life cycle periods, at either 0 to 4 h or 14 to 18 h post merozoite invasion. For Dd2 parasites, cultures were synchronized twice within the same 46-hour life cycle at 0 to 4 h or 14 to 18 h post merozoite invasion.

**Percoll® gradient enrichment.** A 5× RPMI / 25% sorbitol solution was used to dilute pure Percoll® to the required concentration. The Percoll® gradient enrichment was performed by layering 600 µL of 60% Percoll® on top of an equal volume of 90% Percoll® in an Eppendorf tube. Of the pRBC pellet, 200 µL was then layered on the top and the samples spun down in a microfuge at 750 rcf for 10 min.

**Flow cytometry.** In general, trophozoites were isolated using saponin lysis by gently mixing a 0.15% saponin lysis buffer in PBS and a pRBC pellet in approximately a 1:10 ratio. Upon the solution becoming a translucent red color, it was centrifuged at 750 rcf and the supernatant aspirated. The resultant trophozoite pellet was then washed two to three times with 1× PBS and subsequently dissolved in 1× PBS. The trophozoites were then fixed by addition to a counting diluent (0.125% v/v glutaraldehyde and 0.5% v/v DNase I) in a 1:10 ratio and the samples stored at 4 °C for no more than three days prior to running on the flow cytometer. Samples were prepared for flow cytometry by adding together FACS diluent (1× SYBR® Green solution), BD™ Trucount® bead solution (1 mL filtered PBS added to a tube of Trucount® beads and vortexed) and the fixed sample in an 8:1:1 ratio. The samples were then incubated at 37 °C for 30 min and run on a BD™ C6 CSampler flow cytometer at medium fluidics with sample collection limited to 50 µL or 100 µL. All files were exported as FACS files and saved for processing in FlowJo™ (v9 to v10) Software (BD Life Sciences)[3].

*Digestive vacuole studies*

**RBC loading.**[4] RBCs were washed with complete media and then 500 µL added to 1 mL of pre-warmed hypotonic solution (5 mM HEPES, 11 mM glucose and 2 mM NaATP). To produce pHrodo loaded RBCs, to the hypotonic solution 50 µL of 1 mg/mL pHrodo™ dextran stock was added to produce a final concentration of 50 µg/mL. Control RBCs were produced using unmodified hypotonic solution. The hypotonic solutions containing the RBCs were then incubated at 37 °C for 10 min, after which 1 mL of pre-warmed hypertonic solution (300 mM NaCl, 40 mM KCl and 11 mM glucose) was added to create an overall isotonic solution to promote resealing of the RBC membrane. The resealed RBCs were then washed three times with culture medium and used immediately after preparation.

**Incubation and harvesting.** To a 50 mL cell culture flask, 50 µL of a 5% parasitemia pRBC pellet, 200 µL of the resealed RBCs (either containing or not containing pHrodo™) and 13 mL culture medium were added. The flasks were gassed with mixed air and incubated at 37 °C for the required incubation period to achieve certain parasite ages. Following incubation, cultures were spun down and the supernatant aspirated. For NF54 parasites, of the pellet, 5 µL was used to prepare a Giemsa slide blood smear, 200 µL was enriched using Percoll® gradient enrichment and the remainder was saponin lysed, fixed and processed for flow cytometry as described above the following working day. The enriched sample was harvested for confocal imaging at the 60% / 90% interface and washed twice with PBS. For Dd2 parasites, cultures were not enriched, instead 5 µL of the pellet was used to prepare a Giemsa slide blood smear and 50 µL of the pellet was saponin lysed, fixed and processed for flow cytometry. The remainder of the pRBC pellet was used as is for confocal sample preparation. Of the enriched NF54 pellet or Dd2 pRBC pellet, 5 µL was added to 2 mL of Ringer's solution[5] (122.5 mM NaCl, 5.4 mM KCl, 1.2 mM $CaCl_2$, 0.8 mM $MgCl_2$, 11.0 mM glucose, 25.0 mM HEPES and 1.0 mM $NaH_2PO_4$) containing Hoechst stain (1:5000) to produce the final sample for confocal imaging.

**Confocal imaging.** Either single- or four-chambered CellView™ dishes were coated with filtered 0.01% poly-L-lysine and dried at 37 °C prior to use. To the single chamber CellView™ dishes, 500 µL of the confocal sample was added, while 200 µL of the confocal sample was added to the four chamber CellView™ dishes. This was allowed to settle for 5 min before the excess liquid was aspirated. To prevent samples from drying out, fresh Ringer's solution was gently added to the chambers to not disturb the settled cells. Samples were imaged using a Zeiss Confocal Microscope LSM 880, AxioObserver (Airyscan). Settings were established at the during the first experiment and then reused for every subsequent experiment to ensure consistency across different experiments. A C Plan-Apochromat 63× Oil Objective was used with an effective NA of 1.4 and the scan zoom was set to 8.5. The pHrodo™ signal was detected using the red channel (570 to 600 nm band pass filter) while Hoechst was detected using the blue channel (385 to 420 nm band pass filter). The pinhole size for the pHrodo™ channel was 2.31 Airy Units (AU) and the detector gain was set to 700. For Hoechst, the pinhole size was 3.20 AU and the detector gain was set to 850. Using the Z-stack function in Zen Black (Zeiss), Z-stacks of individual parasites were acquired and subsequently subjected to Airyscan processing. The original files in their .czi format were transferred to a storage device and then saved on OneDrive. All samples were imaged within 6 hours of sample collection to ensure sample integrity.

**Image analysis.** The files were processed in Zen Blue Lite (Zeiss) to produce 3D renderings and images for documents. The volume, uptake and concentration analyses were performed using ImageJ.[6, 7] The Fiji (Fiji Is Just ImageJ) package was downloaded and installed from the ImageJ website. The .czi files were imported into ImageJ using the BioFormats Plugin[8] and the color channels separated. The red channel Z-stacks were then thresholded using the built-in RenyiEntropy automatic threshold. The volume analysis was conducted in two ways. In one instance, the "Analyze Particles" (AP) built-in function on ImageJ was used in which the calculations were limited to the threshold and the circularity set to 0.05 to infinity. The resultant area of each region of interest were then summed in Excel and multiplied by the Z-stack spacing to obtain the volume. In the second instance, the 3D Objects Counter Plugin (3D OC)[9] was used to calculate the volume by applying the RenyiEntropy automatic threshold to the Z-stack using the stack histogram and subsequently applying the 3D OC function to the resultant thresholded Z-stack. All the volume data was then collated and compared in Microsoft Excel and analyzed using GraphPad Prism (v8 to v10, GraphPad Software Inc, USA)[10]. To determine the uptake and concentration of the DV, the raw integrated density was obtained for each Z-stack from the file generated by the AP function. The uptake was determined to be the summation of the integrated density of each Z-stack. To calculate the concentration, this value was divided by the volume calculated for that individual parasite.

*PM I and IV protein analysis.*

**Parasite lysate preparation.** The saponin lysis protocol was adopted to isolate the NF54 trophozoites. To the isolated trophozoite pellet, 45 µL filtered PBS was added, the solution was mixed thoroughly and 5 µL of the cell suspension was added to 45 µL of counting diluent. The fixed sample was processed for flow cytometry as described above in 96-well plates and used to calculate the cell concentration. To the remaining 40 µL of the isolated trophozoite cell suspension, 10 µL of 5× sample application buffer (SAB, 5% β-mercaptoethanol solution in 5× SDS running buffer) was added. The sample was mixed thoroughly and heated at 90 °C for 5 min using a heating block. The sample was then stored at -20 °C until further use. All samples needed for the SDS-page were diluted with 1× SAB to a final volume of 10 µL such that 50 000 parasites were contained in the lysate.

**SDS-page.** All back- and spacer plates (casting plates) were washed and cleaned with 70% EtOH prior to use. SDS-page gels consisting of 12% resolving gel (12% acrylamide/bis-acrylamide 37:1, 375 mM Tris pH 8.8, 0.1% w/v SDS, 0.05% v/v TEMED, 0.1% w/v ammonium persulphate) and 5% stacking gel (5% acrylamide/bis-acrylamide 37:1, 250 mM Tris pH 6.8, 0.1% SDS w/v, 0.1% v/v TEMED, 0.1% w/v ammonium persulphate) at 1 mm thickness and containing 15 lanes were hand-cast using a Bio-Rad Mini-Protean 3 casting chamber. Page gels not used immediately were wrapped airtight in cling wrap and stored at 4 °C. Stored page gels were used within 24 hours of casting. A Bio-Rad Mini-Protean Tetra System electrophoresis tank was filled with 1× SDS running buffer (25 mM Tris, 250 mM glycine and 0.5% w/v SDS) and assembled with the page gel. The sample wells were flushed with 1× SDS running buffer prior to loading and to each well 10 µL of parasite lysate or 3 µL NEB Color Prestained Protein Standard Broad Range (P7719S, 10 – 250 kDa) were added according to a predetermined loading pattern. The electrophoresis was run for 45 to 60 min at a constant current of 30 mA per gel and variable voltage. Page gels were removed and immediately processed for the western transfer.

**Western blot preparation.** A PVDF membrane was activated for 15 sec in MeOH and then placed in 1× western transfer buffer (25 mM Tris, 250 mM glycine and 10% v/v MeOH) along with the other components needed for the blotting cassette to equilibrate for 5 min. The blotting cassette was then assembled as per the manufacturer guidelines and placed into the electrophoresis tank which was then filled with 1× western transfer buffer. The transfer was run for 1 hour and 40 min at a constant voltage of 100 V and variable current. Once completed, the PVDF membrane was washed three times with 1× TBST (20 mM Tris pH 8.0, 150 mM NaCl and 0.1% Tween® 20) for 5 min and transferred to blocking buffer (5% w/v skim milk solution in TBST) for 40 min. Once blocked, the membrane was incubated with primary antibody solutions overnight at 4 °C with gentle shaking. Anti-PM I (rabbit, 1: 10 000), anti-PM IV (mouse, 1:10 000) and anti-BiP (rabbit, 1:20 000) were all prepared using blocking buffer. Following incubation with the primary antibodies, the membrane was washed three times with TBST for 5 min and treated with horse radish peroxidase (HRP)-conjugated secondary anti-rabbit (1:3 000) or anti-mouse (1:3 000) antibodies for 1 hour with gentle shaking. The membranes were then washed three times with TBST for 5 min. The Advansta WesternBright® Sirius® detection kit was used for signal detection. Briefly, the PVDF membrane was treated for 2 min with 1 mL of a 1:1 ratio mixture of luminol/enhancer solution and peroxide chemiluminescent detection reagent. Any excess solution was removed, and the membrane secured in cling wrap in a light-proof X-ray cassette. In a dark room, the signal was detected by exposure to X-ray film, which were subsequently developed, fixed, and rinsed with tap water. Once dry, the X-ray films were scanned using an Epson Perfection V800 scanner using Epson Scan Software and stored in the .tiff format.

**Densitometric blot analysis.** Scanned blots were imported into ImageJ and analyzed as previously described.[11] Briefly, .tiff files were converted to grey scale 8-bit images and a background 8-bit image created using the "Subtract Background" function with "Create background – don't subtract" activated and the rolling ball radius set to 50.0 pixels. Using the "Image Calculator", the background image was subtracted from the original 8-bit image to generate a negative image that was then inverted using the "Look Up Tables" (LUT) to generate an image with a white background and dark foreground pixels. The blot was then analyzed using the "Gels" function to produce histograms for each band. These were then quantified using the "Wand" tool and the data acquired as arbitrary area values. Using the "Label Peaks" function, the fractional area of each band was obtained relative to each other. The raw data were then exported into Excel and GraphPad Prism for further analysis.

*Basal levels of the heme-containing species*[12]

**Incubation and harvesting.** NF54 parasites at 4% parasitemia, 2% hematocrit and 2 mL culture volume were incubated in flat-bottomed 12-well plates (Greiner) in an airtight gas chamber flushed with mixed air at 37 °C for variable incubation periods. Given that Dd2 parasites have faster growth rates than NF54, these were prepared at 3% parasitemia, 2% hematocrit and 6 mL culture volume in flat-bottomed 6-well plates (Greiner). Following the required incubation period for harvesting parasites at a given age, excess media supernatant was aspirated, and Giemsa-stained thin smears were prepared and visually analyzed to assess any changes to the parasite population. The pellets were resuspended in 1.8 mL 1× PBS and transferred to deep-well 2.2 mL 96-well plates (Greiner). To this, 125 µL 1% saponin and following gentle shaking for 2 min, the plates were centrifuged for 15 min at 750 rcf and the supernatant aspirated. The isolated trophozoite pellets were washed two to three times with 0.5 mL 1× PBS followed by a 15 min centrifugation at 750 rcf after each wash to remove all erythrocyte hemoglobin. The isolated trophozoites were then resuspended in 100 µL filtered 1× PBS and accurately transferred to a round-bottomed, 96-well 0.5 mL plate (Axygen) referred to as the "stock plate". Of this, 10 µL was transferred to a flat-bottomed 96-well plate (Greiner) containing 190 µL counting diluent. This plate was referred to as the "counting plate". The stock plate and counting plate were then stored at -20 °C and 4 °C, respectively, until further use. The counting plate was then processed for flow cytometry as described above to determine cell count.

**Cellular fractionation.** Following thawing of the stock plate, 100 µL of MilliQ water (m$H_2O$) was added to each well and the plate sonicated for 5 min in an ultrasound bath. To this, 50 µL 0.2 M HEPES buffer (pH 7.5) was added, and the plate centrifuged at 3600 rpm for 20 min. Carefully, without disturbing the pellet, the supernatant was transferred to an adjacent set of wells on the same plate. To the supernatant, 50 µL 4% w/v SDS was added, the plate sonicated for 5 min and incubated at room temperature for 30 min. Following incubation, 50 µL 0.3 M NaCl and 50 µL 25% v/v pyridine (25% v/v in 0.2 M HEPES pH 7.5) in 0.2 M HEPES (pH 7.5) were added and 200 µL of this solution transferred to a flat-bottomed, 96-well plate termed the "reading plate". This fraction corresponded to the hemoglobin fraction.

The pellet was treated with 50 µL $mH_2O$ and 50 µL 4% SDS and resuspended well. The plate was sonicated for 5 min and incubated at room temperature for 30 min. Following incubation, 50 µL each of 0.2 M HEPES (pH 7.5), 0.3 M Na Cl and 25% pyridine were added, and the plate centrifuged at 3600 rpm for 20 min. Without disturbing the delicate pellet, the supernatant was transferred to an adjacent set of well on the same plate. The supernatant was diluted to 400 µL with $mH_2O$ and 200 µL of this solution was transferred to the reading plate. This fraction corresponded to the heme fraction.

The remaining pellet was treated with 50 µL 0.3 M NaOH and 50 µL $mH_2O$, the plate sonicated for 15 min and incubated at room temperature for 30 min. To this, 50 µL each of 0.2 M HEPES (pH 7.5), 0.3 M HCl and 25% pyridine was added, and finally, 150 µL of $mH_2O$. Of this, 200 µL was transferred to the reading plate. The UV-visible spectra of heme as an Fe(III)PPIX-(bis)pyridyl complex was recorded between 400 and 415 nm on a multi-well plate reader (ThermoScientific MultiskanGO). The absorbance maxima of each well were used to calculate the absolute amount and percentage of each heme-containing species in each sample.

**Heme standard curve.** The absolute amount of each heme-containing species was quantified using a heme standard curve. A 100 µg/mL standard solution of hematin (porcine) in 0.3 M NaOH was prepared and 200 µL added to a round-bottomed, 96-well plate (Axygen). Serial dilutions of the standard solution were carried out across the plate using 100 µL 0.3 M NaOH, leaving a blank column for reference. Of each of the following solutions, 50 µL were added to 100 µL of hematin standard: 0.2 M HEPES (pH 7.5), 4% w/v SDS, 0.3 M NaCl, 0.3 M HCl, 25% v/v pyridine in 0.2 M HEPES (pH 7.5) and $mH_2O$. The UV-visible spectra of heme as the Fe(III)PPIX-(bis)pyridyl complex from 400 to 415 nm were recorded using a multi-well plate reader. The standard curve was used to calculate the absolute amount of each heme-containing species as Fe (fg/mL) using the following equation:

$$M_{Fe} = M_H \times \left(\frac{MW_{Fe}}{MW_H}\right)$$

$M_{Fe}$ = mass of Fe
$M_H$ = mass of hematin
$MW_{Fe}$ = molecular mass of Fe
$MW_H$ = molecular mass of hematin
